# NOTE ON RHEIFORMES (AVES: PALAEOGNATHAE) FROM A QUATERNARY CAVE IN THE LAGOA SANTA KARST, EASTERN BRAZIL

**DOI:** 10.64898/2026.09.11.751023

**Authors:** Artur Chahud

**Affiliations:** Institute of Biosciences, University of São Paulo

**Keywords:** Rheidae, *Rhea americana*, Pleistocene, Holocene, Palaeognathae

## Abstract

Rheiformes are a South American lineage of large, flightless palaeognathous birds, currently represented by the genus *Rhea* and the two extant species *R. americana* and *R. pennata*. The fossil record of the group extends back to the Eocene, with its greatest diversification occurring during the Neogene. During the Quaternary, Rheiformes were widely distributed across southern and central South America. The Lagoa Santa region is particularly notable for its rich paleontological and archaeological record. Quaternary records of Rheiformes from caves in this region provide important evidence for understanding the distribution of the group in Brazil, although their chronological context remains uncertain owing to the absence of direct dating. Here, we describe and illustrate two rheiform bones recovered from a cave in the Lagoa Santa karst region. The material comprises a partially complete tibiotarsus and a fragmented tarsometatarsus, interpreted as belonging to a single individual. Comparative anatomical and morphometric analyses support their attribution to *Rhea americana*, based on morphological features and dimensions consistent with adult specimens of the species. This occurrence extends the Quaternary record of Rheiformes in the Lagoa Santa region and contributes to the fossil and subfossil record of *Rhea* in Brazil. The absence of stratigraphic and geochronological control precludes a precise age determination. Although the association with an extinct dasypodid indicates the presence of Late Pleistocene or Early Holocene faunal elements at the locality, a Holocene or even recent age for the rheiform remains cannot be excluded.

## INTRODUCTION

Rheiformes constitute a distinct South American lineage of large, flightless palaeognathous birds, whose extant diversity is restricted to the genus Rhea Brisson, 1760. Only two extant species of Rheiformes are currently recognized: *Rhea americana* Linnaeus, 1758, and *Rhea pennata* d’Orbigny, 1834.

Initially, this clade of birds was included within the group known as “Ratites,” which comprised all flightless birds. However, the traditional concept of “Ratites” as a monophyletic group has been revised and is now considered polyphyletic. It is currently recognized that independent losses of flight occurred repeatedly within the infraclass Palaeognathae, with tinamous, a group that retains the ability to fly, being nested within this broader radiation (Laurin et al., 2012; Yonezawa et al., 2017; Sackton et al., 2019).

The oldest fossil attributed to Rheiformes belongs to the species *Diogenornis fragilis* Alvarenga, 1983, from the Early Eocene of the São José de Itaboraí Basin, in the state of Rio de Janeiro. However, the greatest radiation and diversity of the group occurred during the Neogene, with most species reported from the Miocene and Pliocene of Argentina (Agnolín and Chafrat, 2015; Noriega et al., 2017; Picasso et al., 2022). The earliest record of the genus *Rhea* dates to the Miocene, represented by the species *Rhea mesopotamica* Agnolín and Noriega, 2012.

Quaternary records extend the known distribution of the group to southern and central portions of South America, broadly corresponding to the current distributions of the two extant species, although the latter exhibit marked ecological differences. *Rhea americana* inhabits extensive open landscapes of central and eastern South America, whereas *R. pennata* is mainly associated with Patagonian and high-altitude Andean environments.

The Lagoa Santa region is characterized by extensive karst landscapes and is particularly notable for its exceptional archaeological and paleontological richness. Scientific investigations of faunal assemblages from caves in the region date back to the pioneering studies of Peter Wilhelm Lund (Lund, 1840, 1841, 1842; Holten and Sterll, 2011), whose discoveries contributed decisively to establishing the region as an area of international importance for paleontological research.

The Lagoa Santa region also contains Quaternary records attributed to Rheiformes, recovered from different localities investigated by Peter Lund. Although the age of these specimens has not been directly determined using dating methods, the materials have traditionally been assigned to species currently living in the region.

Between 2001 and 2009, the research project coordinated by Walter Neves, entitled “Origins and Microevolution of Humans in the Americas: A Paleoanthropological Approach,” contributed significantly to expanding knowledge of the archaeological and paleontological record of the Lagoa Santa region. Among the materials recovered during this period was a collection of osteological remains attributed to extinct and extant taxa. These remains were recovered from an unmapped vertical cave informally known as “Abismo Quaternário.” Its precise location is currently unknown; however, it is known to be situated within the same carbonate massif and in the vicinity of Cuvieri Cave, one of the most important vertebrate paleontological sites in the region, whose fauna has been investigated in several studies, particularly those focused on fossil mammals (Chahud and Okumura, 2021a, 2021b; 2023; Chahud et al. 2023a; 2023b; Gomes & Chahud, 2026).

At “Abismo Quaternário,” a tibiotarsus and a tarsometatarsus attributed to Rheiformes were recovered. Thus, the present study aims to describe and taxonomically identify these specimens, as well as to discuss aspects related to the region’s paleoenvironment and associated fauna, and to provide brief comments on the occurrence of Rheiformes in Brazil.

## MATERIALS AND METHODS

The two specimens analyzed in this study were recovered from the vertical cave informally known as “Abismo Quaternário,” which is located in the same carbonate massif as Cuvieri Cave (UTM coordinates 23K 603756 E, 7846105 S), in the municipality of Matozinhos, state of Minas Gerais, eastern Brazil (Figure 1). The locality is situated a few kilometers from Cuvieri Cave within the Lagoa Santa karst region, which is well known for its Quaternary fossil-bearing deposits (Hubbe et al. 2011; Chahud, 2020a; 2020b; 2022; 2026; Ramos and Chahud, 2026).

**Figure 1.**
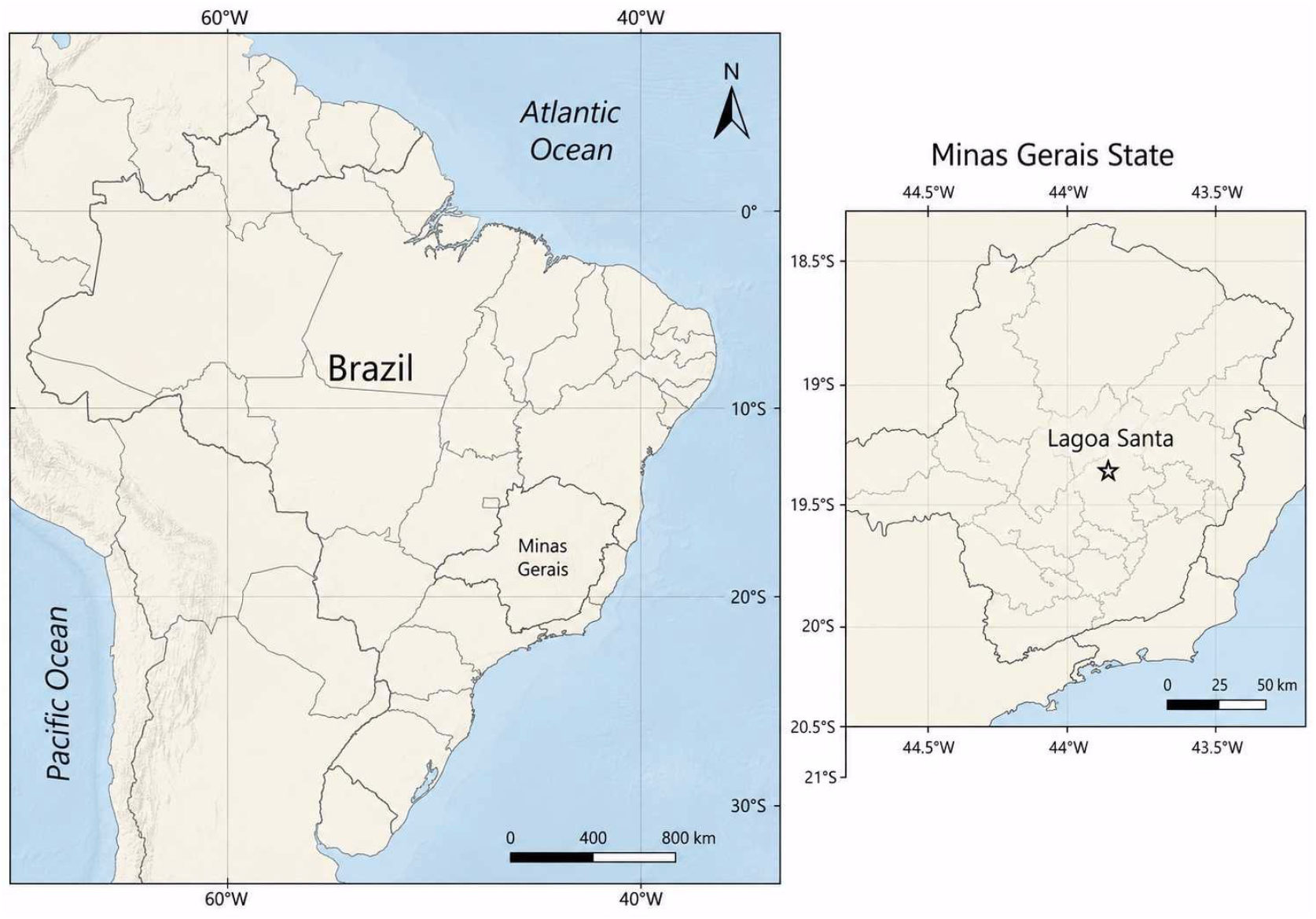
Location of the Lagoa Santa area

Information on the provenance of the material is limited, as there are no records concerning the collection method or the stratigraphic position of the fossils. Both specimens share the same catalog number (AQSE-2), indicating that the specimens were collected together in the cave.

Because the collections made at “Abismo Quaternário” did not follow stratigraphic protocols and it was not possible to date the specimens, the age of the specimens could not be determined precisely. However, the occurrence of an extinct species of Dasypodidae previously recorded from the same locality (Baldavira Coutinho and Chahud, 2026) demonstrates the presence of megafauna elements from the Early Holocene or Pleistocene. Therefore, we adopt a broad chronological assignment to the Quaternary, avoiding more restrictive inferences given the absence of stratigraphic and/or geochronological control.

Taxonomic identification of the specimens and determination of their ontogenetic stage were carried out through anatomical comparisons with previously identified reference material, including a specimen deposited in the zoological collection of the Institute of Biosciences of the University of São Paulo (IB-USP), as well as data described in the specialized literature (Agnolin and Noriega, 2012; Picasso, 2012; Medina et al. 2019; Picasso et al. 2022).

For morphometric measurements, the protocol proposed by Picasso (2012) was adopted. This protocol establishes standard anatomical landmarks for osteological measurements, ensuring the reproducibility of the data in future studies (Figure 2).

**Figure 2.**
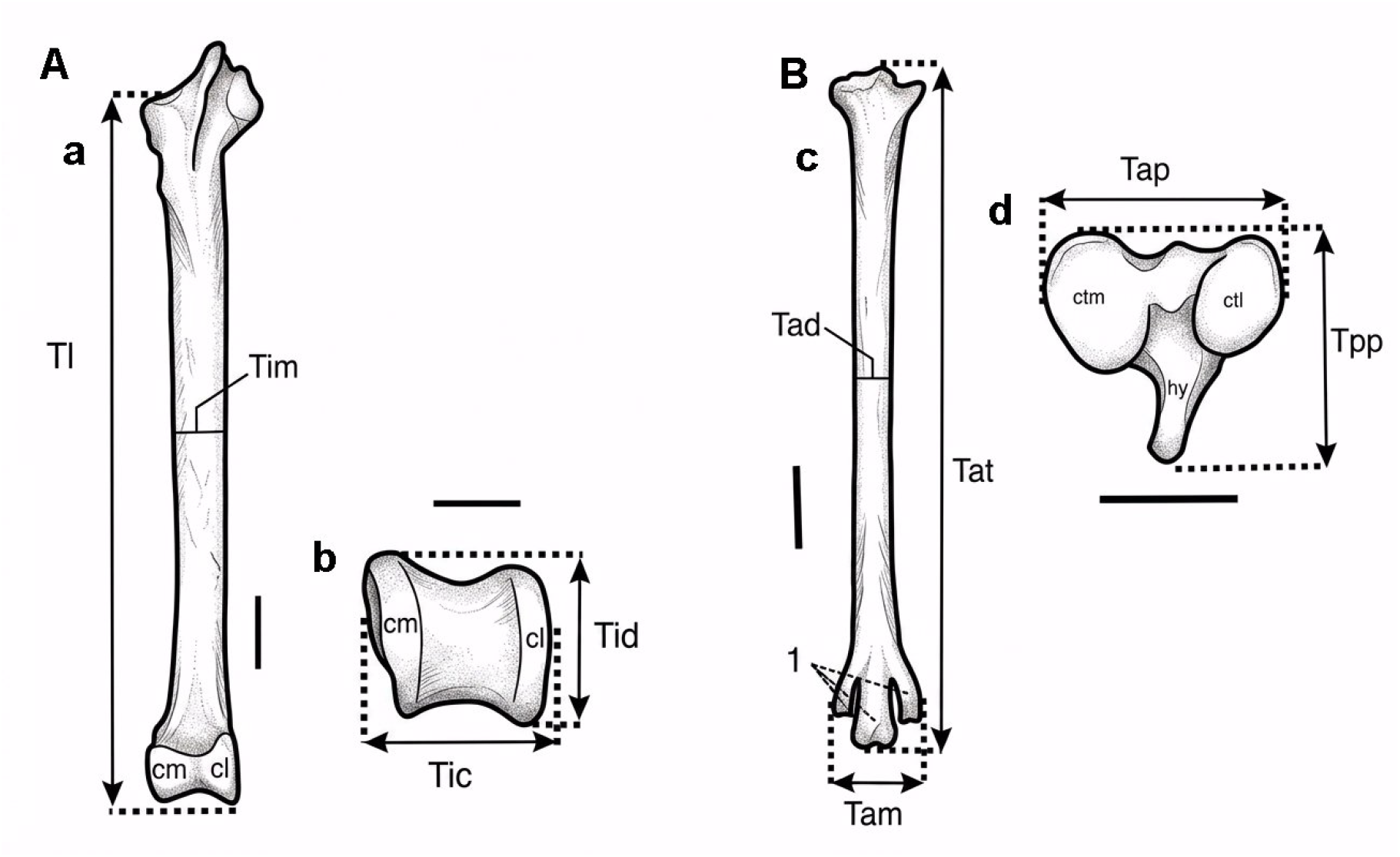
Schematic models of measurements. A) Tibiotarsus: (a) frontal view, (b) detailed view of the distal articular surface; B) Tarsometatarsus: (c) frontal view, (d) detailed view of the proximal articular surface. Abbreviations: cm, *condylus medialis*; cl, *condylus lateralis*; ctm, *cotyla medialis*; ctl, *cotyla lateralis*; hy, hypotarsus. 1, trochlea metatarsii, II, III and IV (from left to right); Tl: tibiotarsus length; Tim: latero-medial diameter of *corpus tibialis*; Tid: antero-posterior width of the distal; Tic: latero-medial width of the distal; Tat: total length; Tad: latero-medial diameter of corpus tibiotarsi; Tap: latero-medial width of the proximal; Tpp: antero-posterior width of the proximal; Tam: latero-medial width of the distal. Scale bar: 10 mm. Adapted from Picasso (2012).

### QUATERNARY RHEIFORMES OF THE BRAZILIAN TERRITORY

The Quaternary records of Rheiformes in Brazil are represented by the genus *Rhea*. In Brazilian territory, the genus is identified primarily by the species *Rhea americana*, in addition to specimens determined only as *Rhea* sp. and cf. *Rhea* sp., reflecting different degrees of confidence in taxonomic identification (Nascimento and Silveira, 2024). The occurrences are distributed across several Brazilian states, including Ceará, Goiás, Minas Gerais, Mato Grosso do Sul, Pernambuco, Rio Grande do Sul, São Paulo, and Piauí (Faure et al. 2010; Nascimento and Silveira, 2024).

Among the most relevant records are those from caves, rock shelters, natural tanks, and other Quaternary deposits. In the Lagoa Santa region, Minas Gerais, records associated with caves and rock shelters are particularly noteworthy, constituting one of the main localities where Rheiformes occur in the Brazilian Quaternary. (Lund, 1840; 1841; 1842; Winge, 1887; Nascimento and Silveira, 2020; 2024). In Piauí, Toca do Serrote das Moendas, located in the Serra da Capivara area, contains material attributed to *Rhea* sp., represented by a distal end of a tarsometatarsus (Guidon et al. 2009; Faure et al. 2010; Nascimento and Silveira, 2024). This specimen was previously identified as *Rhea fossilis* by Faure et al. (2010), but it is now classified more conservatively (Nascimento and Silveira, 2024).

In Ceará, records from the Itapipoca region also provide evidence of the presence of Rheidae in Quaternary deposits, including material associated with natural tanks (Ximenes 2009; Waldherr et al. 2017, 2023; Nascimento and Silveira 2024). The record of *Rhea americana* from Ceará is represented by skeletal material that may correspond to a juvenile or subadult individual (Ximenes, 2009; Waldherr et al. 2017, 2023; Costa et al. 2024; Nascimento and Silveira, 2024).

In addition to these sites, *Rhea americana* has been recorded in several other states, demonstrating a broad geographic distribution during the Brazilian Quaternary. The recovered materials include various bones, fragments of long bones, tarsometatarsi, and eggshell fragments, as well as remains associated with archaeological and subfossil contexts (Nascimento and Silveira, 2020; 2024).

### SYSTEMATIC PALEONTOLOGY

Class Aves Linnaeus, 1758

Infraclass Palaeognathae Pycraft, 1900

Order Rheiformes Forbes, 1884

Family Rheidae Bonaparte, 1849

Genus *Rhea* Brisson, 1760

Species *Rhea americana* (Linnaeus, 1758)

Figure 3

**Figure 3.**
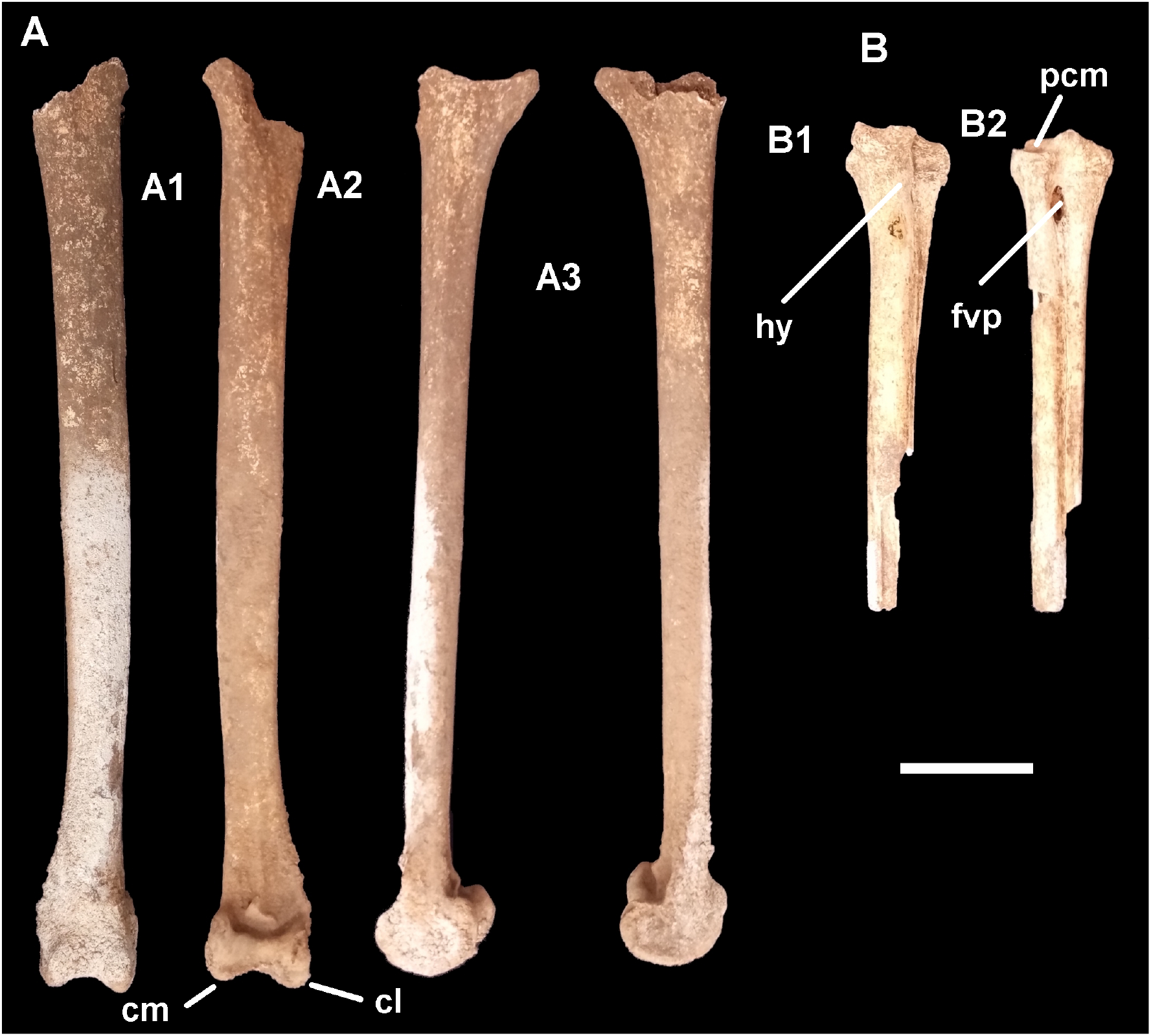
Bone parts of *Rhea americana* found in the “Abismo Quaternário”. A) Tibiotarsus, A1) Posterior view; A2) Anterior view; A3) Lateral views; B) Tarsometatarsus, B1) Plantar view; B2) Dorsal view. **Abbreviations:** hy: hypotarsus, fvp: foramen vascular proximal, pcm: prominence of the medial cotyle. Bar scale: 50 mm.

### Locality

Lagoa Santa Karst, Minas Gerais, Brazil, “Abismo Quaternário”.

### Material

One partially complete tibiotarsus and one incomplete tarsometatarsus, attributed to the same individual.

### Age

Quaternary, associated with regional deposits spanning the Pleistocene–Holocene.

### Remarks

The specimens from the “Abismo Quaternário” consist of a tibiotarsus and a tarsometatarsus whose proportions are consistent with those of an adult *Rhea americana* (Table 1). The tibiotarsus exhibits an elongated, predominantly straight shaft with relatively uniform dimensions along its midshaft region. The bone gradually expands toward both ends; the proximal region is fragmented, preventing measurement of the specimen’s maximum length, whereas the distal region preserves the condyles.

**Table 1.** Main measurements obtained for the tibiotarsus and tarsometatarsus from the “Abismo Quaternário”. **Abbreviations:** Tim: latero-medial diameter of the corpus tibialis, Tid: antero-posterior width of the distal end taken between extreme points, Tic: latero-medial width of the distal end taken between extreme points of both condyles of tibiotarsi, Tad: latero-medial diameter of corpus tibiotarsi, Tap: latero-medial width of the proximal end taken between extreme points of both cotyla, Tpp: antero-posterior width of the proximal end, taken from the most extreme point of the hypotarsus to the most extreme point of the anterior edge. Range based on known adult specimens and on data from Picasso (2012) and Medina et al.(2019).

| **Tibiotarsus** |  |  |  |
| --- | --- | --- | --- |
|  | Tim | Tid | Tic |
| Range (N=8) | 31.63 – 20.74 | 42.45 – 34.05 | 39.99 – 33.82 |
| **Specimen AQSE-2** | **24.8** | **37.7** | **37.4** |
| **Tarsometatarsus** |  |  |  |
|  | Tad | Tap | Tpp |
| Range (N=11) | 19.7 – 14.46 | 43.05 – 36.09 | 38.85 – 34.08 |
| **Specimen AQSE-2** | **17.4** | **39.26** | **37.8** |

In the proximal portion of the shaft, the lateral surface of the tibiotarsus exhibits the fibular crest (*crista fibularis*), which is fragmented (Figure A2) at its extremity; nevertheless, its morphology was sufficient to establish that the element was from the right side. The same inference can be applied to the tarsometatarsus since the piece was originally articulated with the tibiotarsus.

The distal end of the tibiotarsus is expanded and terminates in two main articular prominences, the medial condyle (*condylus medialis*) and the lateral condyle (*condylus lateralis*), separated by the intercondylar region (Figures A1 and A2), which delimits the articular surfaces and participates in the articulation with the tarsometatarsus; both structures are preserved. The condylar surfaces exhibit a rounded outline and continuity with the shaft, forming the region responsible for transmitting forces to the tarsometatarsus.

The tarsometatarsus preserves only the proximal portion and part of the shaft. The proximal articular region is markedly expanded relative to the shaft, corresponding to the articular area that receives the condyles of the tibiotarsus (Figure B). The preserved portion of the shaft is elongated, narrow, and relatively straight, consistent with the pattern observed in representatives of the genus *Rhea*.

The hypotarsus (Figure B1), although worn, is visible, as are the proximal vascular foramen (*foramina vascularia proximalia*) and the dorsal infracotylar fossa (*fossa infracotylaris dorsalis*), both of which are clearly distinguishable (Figures B1 and B2). Comparison with extant specimens indicates that the size and proportions of the tibiotarsus and tarsometatarsus are consistent with their attribution to a single adult individual.

## Discussion

The earliest records of *Rhea americana* in caves from the Lagoa Santa region date back to studies conducted by Peter Wilhelm Lund, mainly between 1835 and 1844. The occurrences documented during this period come predominantly from Lapa da Anna Felícia, Lapa da Anta I, and Lapa da Escrivânia I, with additional material possibly grouped from the caves of Coxos, Ossinhos, and Serra das Abelhas (Lund, 1840; 1841; 1842; Winge, 1887; Nascimento and Silveira, 2024).

Some of this material was initially attributed to other large-bodied birds or to species later considered invalid. At present, however, its identification as Rheiformes is considered reliable, with *Rhea americana* being the only species recognized in Brazil to date.

Other Quaternary taxa from Argentina, previously described as extinct species of *Rhea*, such as *R. anchorenensis, R. pampeana, R. fossilis*, and *R. subpampeana*, are currently treated as synonyms of *Rhea americana* and are no longer considered valid names (Picasso, 2016; Picasso and Mosto, 2016; Picasso et al., 2022). The only valid extinct species of the genus is *Rhea mesopotamica*, from the Miocene of Argentina, represented exclusively by the distal portion of tarsometatarsi from adult and subadult individuals, which prevents direct comparisons with the specimen described herein.

No diagnostic features were preserved that would allow the examined material to be differentiated from other fossil or extant species and subspecies of the genus *Rhea*. Nevertheless, the dimensions of the specimens are compatible with those of adult individuals of *Rhea americana*, exceeding those of the other extant species of the genus and being consistent with measurements observed in modern populations from the Lagoa Santa region. It should be emphasized that the specimens could not be dated, and therefore a Holocene or even recent origin cannot be ruled out. Accordingly, the attribution to *Rhea americana* is considered the most appropriate for the specimens presented herein.

### ASSOCIATED FAUNA AND THE QUATERNARY RECORD

Argentina has an exceptionally rich and well-studied Cenozoic fossil record, whereas other parts of South America remain comparatively undersampled, including Brazil. Therefore, the current geographic distribution of fossil and subfossil records does not reflect the actual occurrence of these taxa during the South American Quaternary. Nevertheless, Argentina provides the only absolute dates currently available for the genus *Rhea*. The oldest of these records, attributed to *Rhea americana* (initially assigned to *R. anchorenensis*), dates to the Ensenadan (Early to Middle Pleistocene) and is the only one that does not fall within the Pleistocene–Holocene transition (Picasso et al. 2022).

The occurrence of *Rhea americana* of the Early Pleistocene is consistent with the persistence of the species throughout much of the Quaternary.

Rheiformes are large cursorial birds that inhabit open and semi-open environments and probably maintained this ecological association throughout the Quaternary.

When examining the specimens found in Lagoa Santa and comparing them with the specimens described by Lund, it can be observed that, during the Quaternary, the genus *Rhea* occurred in association with remains of large herbivorous mammals, camelids, cervids, equids, xenarthrans (ground sloths, glyptodonts, and large armadillos), and carnivores. These associations vary considerably among localities and should not be interpreted as representing a single type of community because, to date, no specimen of *Rhea americana* from Brazil has been successfully dated or adequately placed within a stratigraphic deposit. Therefore, some or even all of these specimens could be Holocene or recent in age. The same limitation applies to the specimens from “Abismo Quaternário”, even though these have been associated with an extinct species of Dasypodidae (Baldavira Coutinho and Chahud, 2026).

The archaeological record in Brazil has received little attention in recent research, with studies largely reiterating Lund’s observations within taxonomic works. At archaeological sites, rock paintings, bone remains used as artifacts, food remains, and eggshells have been recorded (Guidon et al., 2009; Nascimento and Silveira, 2024), demonstrating that humans interacted with Rheiformes in several ways throughout much of the Holocene.

### FINAL CONSIDERATIONS

The material analyzed herein, consisting of a partially complete tibiotarsus and a fragmented tarsometatarsus from “Abismo Quaternário,” expands the fossil record of Rheiformes in the Lagoa Santa karst region and in the Brazilian Quaternary. The specimens exhibit anatomical features and proportions compatible with those of an adult individual of *Rhea americana*. This identification is supported primarily by the morphology of the preserved elements and by the measurements obtained, which fall within the ranges observed in adult specimens of the species and further support their attribution to a single individual, given the proportional compatibility between the dimensions of the tibiotarsus and tarsometatarsus.

The absence of stratigraphic control and direct dating, together with the lack of precise information regarding the collection method, prevents the establishment of a more precise age for the specimens. Although the association with an extinct species of Dasypodidae indicates the presence of elements attributable to the Late Pleistocene or Early Holocene in “Abismo Quaternário”, a Holocene or even recent origin for the *Rhea* material cannot be excluded. Thus, a broad Quaternary assignment is the most appropriate interpretation given the available evidence.

The presence of the genus *Rhea* further supports the existence of open and semi-open environments in the Lagoa Santa region during the Quaternary, which did not differ substantially from those occurring in the region today.

